# High-amplitude co-fluctuations in cortical activity drive functional connectivity

**DOI:** 10.1101/800045

**Authors:** Farnaz Zamani Esfahlani, Youngheun Jo, Joshua Faskowitz, Lisa Byrge, Daniel P. Kennedy, Olaf Sporns, Richard F. Betzel

## Abstract

Resting-state functional connectivity is used throughout neuroscience to study brain organization and to generate biomarkers of development, disease, and cognition. The processes that give rise to correlated activity are, however, poorly understood. Here, we decompose resting-state functional connectivity using a “temporal unwrapping” procedure to assess the contributions of moment-to-moment activity co-fluctuations to the overall connectivity pattern. This approach temporally resolves functional connectivity at a timescale of single frames, which enables us to make direct comparisons of co-fluctuations of network organization with fluctuations in the BOLD time series. We show that, surprisingly, only a small fraction of frames exhibiting the strongest co-fluctuation amplitude are required to explain a significant fraction of variance in the overall pattern of connection weights as well as the network’s modular structure. These frames coincide with frames of high BOLD activity amplitude, corresponding to activity patterns that are remarkably consistent across individuals and identify fluctuations in default mode and control network activity as the primary driver of resting-state functional connectivity. Finally, we demonstrate that co-fluctuation amplitude synchronizes across subjects during movie-watching and that high-amplitude frames carry detailed information about individual subjects (whereas low-amplitude frames carry little). Our approach reveals fine-scale temporal structure of resting-state functional connectivity, and discloses that frame-wise contributions vary across time. These observations illuminate the relation of brain activity to functional connectivity and open a number of new directions for future research.

## INTRODUCTION

Resting-state functional connectivity (rsFC) refers to the correlation structure of fMRI BOLD activity, usually estimated over the course of an entire scan session [1, 2]. Inter-individual differences in rsFC have been linked to variation in biological age [3, 4], cognitive state [5], and clinical status [6]. Other studies have emphasized the dynamic nature of rsFC, using sliding window techniques to generate temporally blurred estimates of rsFC across time [7–9] and linking changes in network architecture to behavior [10, 11] and phenotypes [12, 13].

Despite intense interest and widespread application, the processes that underpin and shape rsFC are not fully understood. For instance, how do moment-to-moment fluctuations in connectivity contribute to the pattern of rsFC estimated over longer timescales? How are changes in connectivity supported by instantaneous fluctuations in brain activity? While approaches like innovation-driven co-activity patterns (iCAPs) [14, 15], state-based models [16], and sliding-window analyses [17] have provided insight into the dynamics of either activity or connectivity, they generally are less well suited to provide insight into the relations between domains.

Here, we address these questions using a novel approach for modeling instantaneous co-fluctuations in rsFC [18]. We find that at rest co-fluctuations are “bursty” and occur intermittently as part of whole-brain co-fluctuation “events” that are uncorrelated with respiration, cardiac cycle, and in-scanner motion. We then show that rsFC estimated using only event frames is highly correlated with rsFC estimated over the entire scan session, indicating that rsFC and its system-level organization are driven by co-fluctuations during relatively few frames. We then show that events are underpinned by the activation of a particular spatial mode of brain activity in which default mode and control networks anticorrelated with sensorimotor and attentional systems. We then present two careful examinations of “events.” First, we demonstrate that event time series synchronize across subjects during movie-watching and, second, we show that subjects’ “fingerprints” are enhanced during event frames compared to non-event frames.

## RESULTS

The strength of rsFC between two brain regions can be quantified as the Pearson correlation of their fMRI BOLD time series, which is calculated (after z-scoring) as the mean value of their element-wise product [19]. By omitting the averaging step, we can “temporally unwrap” the correlation measure, which results in a new set of time series – one for every pair of brain regions (network edges) – whose elements represent the magnitude of co-fluctuation between those regions resolved at every moment in time (Fig. 1a). These edge time series can be analyzed directly to pinpoint both the magnitude and timing of co-fluctuations between pairs of brain regions.

**FIG. 1.**
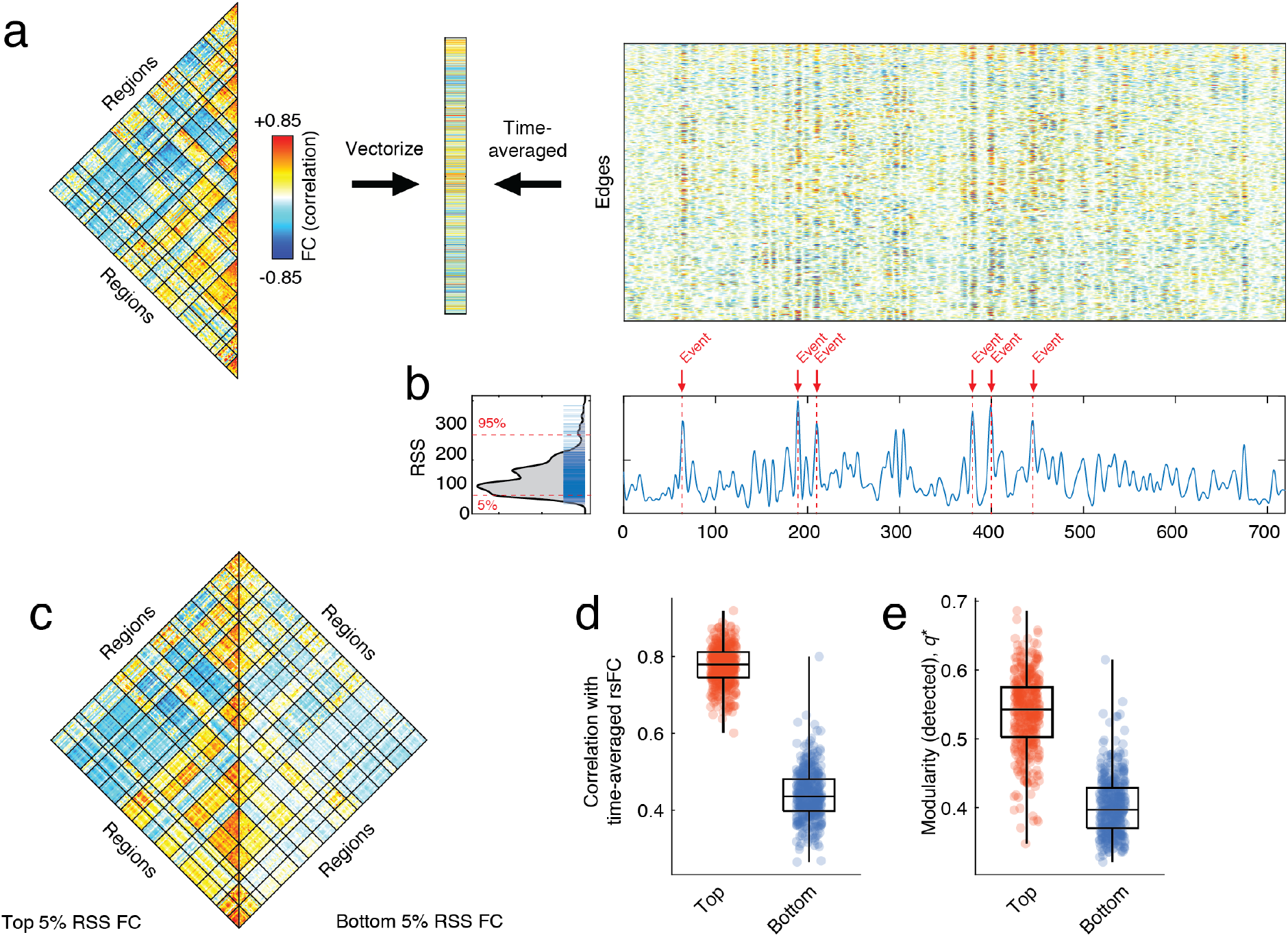
Co-fluctuation time series reveal event structure of resting-state functional connectivity. (*a*) We use a “temporal unwrapping” of the Pearson correlation to generate co-fluctuation time series for every pair of brain regions (edges). The elements of the co-fluctuation time series are the element-wise products of z-scored regional BOLD time series that, when averaged across time, yield vectors that are exactly equal to the Pearson correlation coefficient and can be rearranged to create an resting-state functional connectivity matrix. *(b)* We find that the co-fluctuation time series contains moments in time where many edges collectively co-fluctuate. We can identify these moments by calculating the root sum square across all co-fluctuation time series and plotting this value as a function of time. We consider high-amplitude values as potential “events”. In panel b we label moments in time corresponding to particularly high amplitude co-fluctuations that would be classified as events. The distribution of edge co-fluctuation amplitude is heavy tailed. We wanted to assess the contribution of events and non-events to the overall pattern of functional connectivity. To do this, we extracted the top and bottom 5% of all time points (ordered by co-fluctuation amplitude) and estimated functional connectivity from those points alone. (c) Average functional connectivity across 100 subjects using top 5% (left) and bottom 5% (right). *(d*) In general, the networks estimated using the top 5% of time points were much more similar to traditional functional connectivity than those estimated using the bottom 5% of time points. (e) We performed a similar comparison of network modularity using networks reconstructed using top and bottom 5% frames.

In the first two subsections, we analyze co-fluctuation time series constructed from functional imaging data acquired as part of the Human Connnectome Project [20] (see **Materials and Methods** for details). All results reported in those sections generated using these data; we replicate these findings using a second dataset [21], with results reported in the **Supplementary Material**. In the third and fourth sub-sections, we analyze an independently acquired movie-watching dataset [22] and data from the Midnight Scan Club [21], respectively.

### rsFC is driven by short-lived and high-amplitude co-fluctuation events

When estimated using Pearson correlation, rsFC is expressed as a normalized and time-averaged (over the entire scan session) measure of how strongly the activity of two brain regions co-fluctuates. While past studies have used sliding window methods to generate estimates of moment-to-moment fluctuations in rsFC [8, 9], the use of a windowing procedure results in a temporally blurred estimate of rsFC. This restricts the time scale of observations regarding dynamic changes in functional connectivity to the width of the time window, generally on the order of dozens of frames (1 minute of real time). Here, we address this limitation using co-fluctuation times series, which allow us to accurately assess contributions made to rsFC by single frames without the necessity of a sliding window.

When analyzed across the whole brain, we find that edge time series exhibit “bursty” behavior, such that the amplitude of co-fluctuations (quantified by computing the root sum square; RSS) moves around a mean value, but is punctuated by brief, intermittent, and disproportionately large fluctuations, which we refer to as “events” (Fig. 1b). These events are not directly related to cardiac and respiratory cycles, in-scanner head motion (Fig. S1), and spectral properties of fMRI BOLD time series (Fig. S2), and appear aperiodic with heavy-tailed distributions of event size, event durations, and inter-event intervals (Fig. S3).

To better understand how instantaneous co-fluctuations contribute to whole-brain rsFC, we isolated high-amplitude “events” and compared them with low-amplitude episodes (top and bottom 5% in terms of co-fluctuation amplitude; 60 frames for HCP; see Fig. S4 for comparisons at other percentiles). We then estimated rsFC separately for each category, using only fMRI BOLD data corresponding to those time points and compared the resulting networks. First, we found that connection weights were significantly stronger during events than non-events (within-sample t-test; *p* < 10^-15^; Fig. 1c). Next, we calculated the similarity of rsFC estimated during events and low-amplitude episodes with respect to time-averaged rsFC estimated using the full time series. We found that the event networks were highly correlated with rsFC *(r* = 0.81 ± 0.05) while the non-event networks were much less correlated *(r* = 0.54 ± 0.07) and that these differences were highly significant *(t*-test, *p* < 10^-15^; Fig. 1d). We also performed an analogous comparison of network modularity [23], an index that can be interpreted as a measure of how segregated a network’s systems are from one another. As before, we found that modularity was greater in the event networks *(q* = 0.51 ± 0.06) compared to the non-event networks (q = 0.37 ± 0.05) (*t*-test, p < 10^-15^; Fig. 1e).

In the supplement we show similar results in a second dataset (Fig. S5). We also demonstrate that these effects persist with highly conservative motion censoring (Fig. S6), when using an alternative strategy for estimating networks from the top and bottom 5% time points (Fig. S7), and when comparing against a null model that preserves the temporal structure of events while sampling frames randomly from the entire time series (Fig. S8).

Collectively, these results suggest that rsFC, estimated over long time scales, is driven by a small number of brief, intermittent, and high-amplitude co-fluctuations. The network structure over these points in time contributes disproportionately to the overall modularity and system-level organization of cerebral cortex, as estimated from long-time averages of rsFC. In contrast, low-amplitude co-fluctuations are only weakly related to the overall pattern of rsFC and correspond to less modular architectures.

### FC events are driven by fluctuations of task-positive/task-negative mode of brain activity

In the previous section we demonstrated that time-averaged rsFC can be explained by high-amplitude co-fluctuations that occur during a relatively small number of frames. It remains unclear, however, whether co-fluctuation events are underpinned by a specific pattern of brain activity or whether they reflect contributions from multiple distinct patterns. Here, we address this question directly, by investigating the patterns of brain activity that occur at the same time as events.

As a first point of comparison, we calculated the RSS of both the co-fluctuation time series as well as the z-scored fMRI BOLD time series. We found that, across subjects, these time series were highly correlated (r = 0.97), indicating that co-fluctuation events have an almost one-to-one correspondence with high-amplitude BOLD fluctuations (Fig. 2a). This relationship is expected; because co-fluctuations are calculated as products of z-scored regional activity, their amplitudes will necessarily be correlated with one another.

**FIG. 2.**
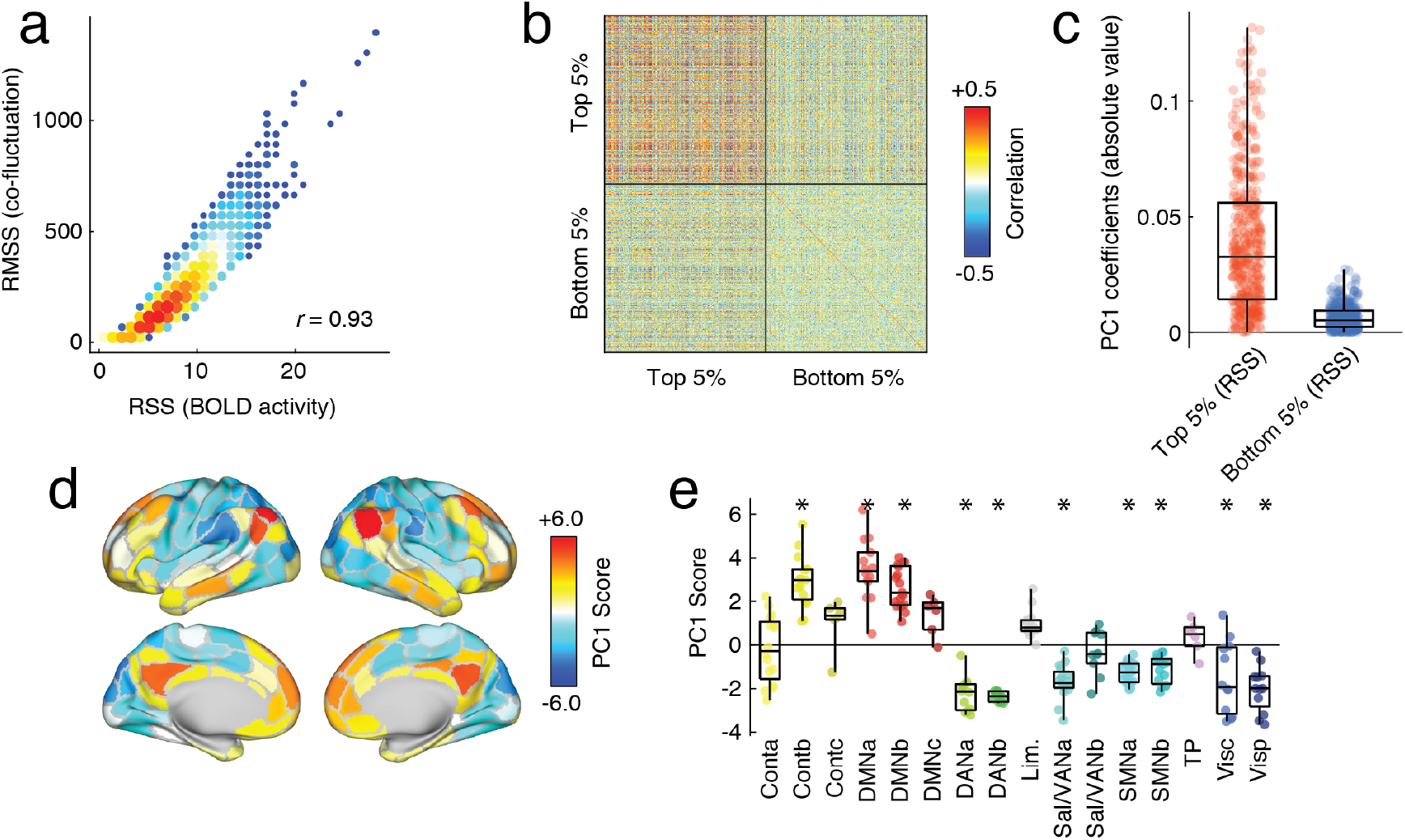
Relationship of network co-fluctuations with BOLD fluctuations. In the previous section we demonstrated that resting-state functional connectivity could be explained on the basis of relatively few frames during which high-amplitude co-fluctuations occurred. Here, we relate those co-fluctuation frames to BOLD activity fluctuations. We first calculate the root sum square amplitude of BOLD activity at each time point and compare that to the amplitude of co-fluctuations. (*a*) Pooling data from across subjects, we find that these two variables are highly correlated. (*b*) To investigate this relationship further, we extract mean activity patterns for each subject and for each scan during the top and bottom 5% time points, indexed according to co-fluctuation amplitude. Here, we show the correlation matrix of those activity vectors. (*c*) We then performed a principal component analysis of this correlation matrix and found that absolute value of coefficients for the first component (PC1) were greater for the top 5% than the bottom 5%, and (*d*, *e*) the PC1 score corresponded to activity patterns that emphasized correlated fluctuations of default mode and control networks that were weakly or anti-correlated with fluctuations elsewhere in the brain. Asterisks indicate systems whose mean PC1 score was significantly greater (more positive or negative) than expected by chance (permutation test; FDR fixed at 5%; *_padjusted_* = 0.018). These observations suggest that co-fluctuation events, which drive resting-state functional connectivity, are underpinned by instantaneous activation and deactivation of default mode and control network areas.

Given that fluctuations in BOLD activity are greater during events than non-events, we asked whether they formed a consistent and recognizable pattern of activity. To address this question, we calculated the mean activity pattern for each subject during their events and nonevents and computed between-subject and between-scan similarity (Fig. 2b). In general, activity during events was more correlated across subjects compared to the activity patterns during non-events *(t*-test, p < 10^-15^). To better understand what was driving these correlations, we performed a principal components analysis of the activity patterns during events and non-events, aggregated over all subjects and scans. We focused on the first principal component (PC1), which explained 26% of total variance. The coefficients for PC1 were, on average, much greater for events than non-events (t-test, p < 10^-15^; Fig. 2c), indicating that PC1 was descriptive of activity patterns during events but less so for non-events. We then mapped component scores for PC1 onto the cortical surface and found that PC1 corresponded to a mode of activity that delineates regions in default mode and control networks from sensorimotor and attentional networks (Fig. 2d,e). We replicated these results in a second dataset (see Fig. S9).

These results suggest that underlying co-fluctuation events is a mode of brain activity whose spatial pattern resembles the traditional task-positive/task-negative division of the brain [24]. This pattern of activity is similar across individuals, suggesting a conserved mechanism by which rsFC emerges from brain activity. These observations suggest a fundamental link between distinct patterns of brain activity and connectivity while further clarifying the origins of co-fluctuation “events.”

### Intersubject synchrony of whole-brain co-fluctuation amplitude during passive movie-watching

In the previous sections we showed that rsFC can be viewed as an average of time-varying co-fluctuations. We also showed that time-averaged rsFC is disproportionately impacted by “event” frames that are, themselves, underpinned by a specific mode of brain activity and were not clearly related to motion or physiological artifacts. What, then, is the purpose of ‘events”? Are they random co-fluctuations or are they related to fluctuations in an individual’s brain/cognitive state? To address these questions, we explored the co-fluctuation time series for a cohort of 29 subjects that were scanned multiple times at rest and while passively viewing complex, naturalistic stimuli (movies) [22].

Specifically, we computed edge time series for all subjects and scans for both conditions. From these edge time series, we estimated the co-fluctuation amplitude across all node pairs. For a given scan, this procedure results in 29 time series (one per subject) of identical length. We found that co-fluctuation time series were correlated across subjects during movie-watching (Fig, 3a,b) but uncorrelated during rest (Fig. 3c,d). We directly compared the distributions of inter-subject correlations between conditions (all scans from the same condition pooled together), discovering that, as expected, the mean intersubject correlation was greater during movie-watching than at rest (permutation test, p < 0.05; Fig. 3e). Importantly, we found no difference between conditions for the overall amplitude of RSS values (permutation test, p = 0.07; Fig. 3f).

**FIG. 3.**
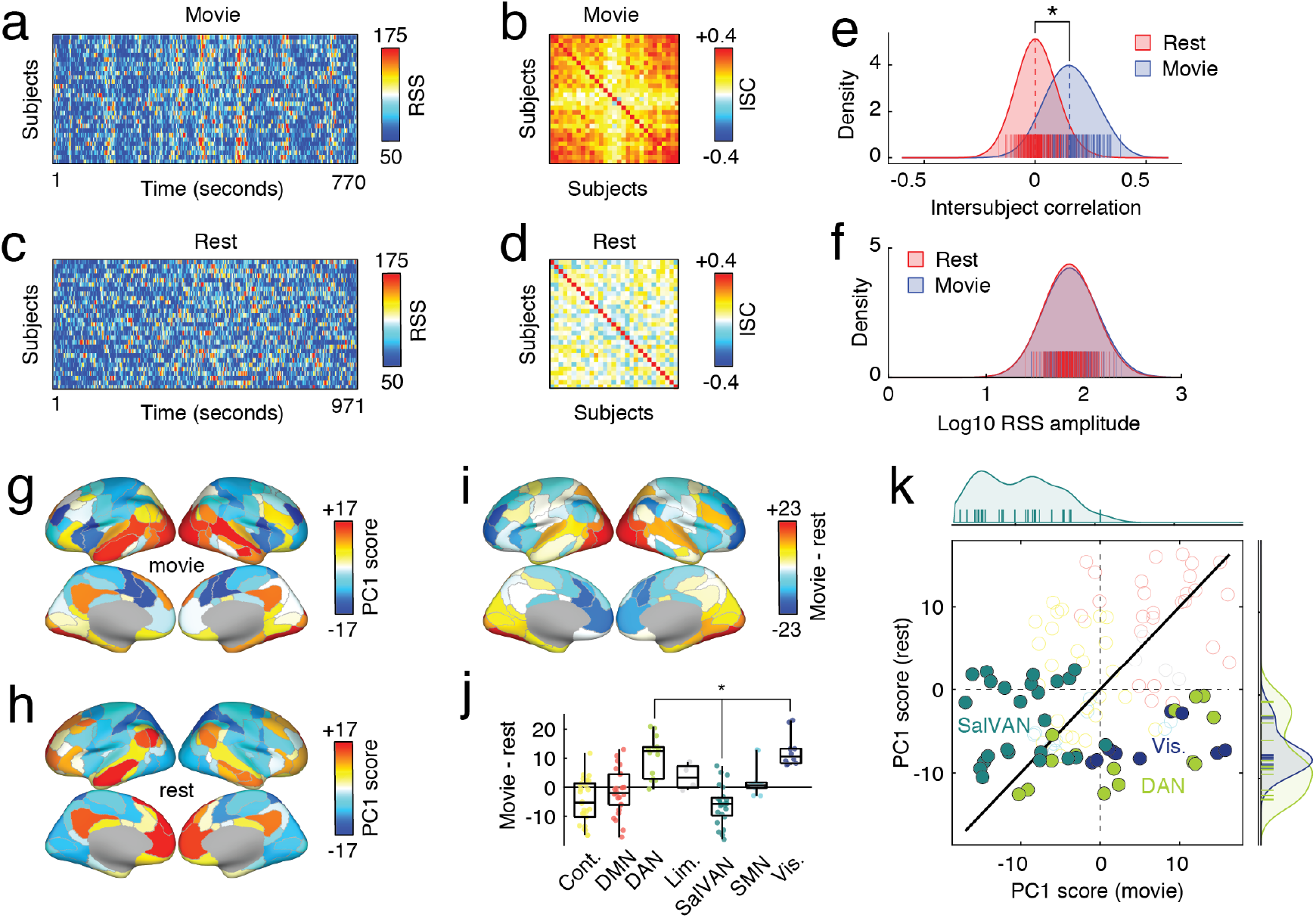
Whole-brain co-fluctuation amplitude synchronizes during passive movie-watching. We compared co-fluctuation time series during resting-state and movie-watching. For both conditions, we computed co-fluctuation time series for 29 subjects. We show those time series in panels *a* (movie) and *c* (rest). We find that when subjects watch movies, their co-fluctuation time series are synchronous, presumably due to the shared audiovisual stimulus. At rest, co-fluctuation time series are asynchronous. We demonstrate this synchrony by computing the inter-subject correlation matrix of subjects’ co-fluctuation time series. We show matrices for movie-watching and rest in panels *b* and *d*, respectively. By comparing the elements of these matrices, we demonstrate statistically that movie-watching leads to increased inter-subject correlations. We show the distributions in panel *e*. We find, however, that the overall amplitude of fluctuations (root sum square; RSS) is not statistically different from one condition to the other (see panel *f*). To further contrast these two conditions, we repeated the analysis from the previous section to identify modes of brain activity that underpins high-amplitude frames. We find that the resting mode recapitulates the topographic distribution reported in the previous section *(h*), emphasizing a task-positive/task-negative division. During movie-watching, however, the mode of activity emphasizes contributions of visual and dorsal attention networks *(g*). In panels *i-k,* we compare rest and movie-watching modes of activity more directly. Panel *i* depicts the regionwise differences in modes, *j* groups those differences by system, and k presents them as a scatterplot, highlighting differences associated with visual, dorsal attention, and salience/ventral attention networks.

Next, we explored differences between movie-watching and rest in terms of brain activity patterns during events and non-events. Our exploration consisted of two analyses. First, and as in the previous section, we extracted activity patterns during events and non-events (top and bottom 5% by co-fluctuation amplitude) separately for the movie-watching resting conditions. We then performed PCA on these matrices and retained the top PC score for each condition. Interestingly, these PC scores exhibited distinct topography; the movie-watching PC (Fig. 3g) emphasized activity in visual and dorsal attention networks, whereas the resting PC (Fig. 3h) recapitulated the pattern shown in the previous section, emphasizing a task-positive/task-negative mode of activity. To directly compare these two patterns, we computed their element-wise (region-wise) difference and grouped these differences by system (Fig. 3i,j). As expected, we found statistically significant differences in the dorsal attention and visual systems (movie > rest; false discovery rate fixed at *q* = 0.05) and salience/ventral attention system (movie < rest). These differences are further evident when we plot the PCs against one another, revealing that these systems deviate from the identity line (Fig. 3k).

We note that another strategy for comparing movie-watching and resting state conditions is to analyze them simultaneously by concatenating event activity patterns from both conditions into a single matrix and jointly decomposing that matrix using PCA. This procedure results in modes of event activity that are *shared* across both conditions. Here, we retain both the first and second PCs (Fig. S10a,c), whose spatial topography is similar to what we show in Fig. 3g,h. As expected, we find differences between the two maps in terms of their PC coefficients, with movie-watching frames loading more strongly onto the first map (permutation test, p < 0.05; Fig. S10b) and resting frames loading more strongly onto the second (permutation test, p < 0.05; Fig. S10d).

Viewed collectively, these results complement our previous findings that co-fluctuation time series are not clearly related to motion or physiological artifacts. Importantly, we demonstrate that subjects’ co-fluctuation time series synchronize when jointly presented with complex, time-varying, and naturalistic stimuli. This observation, combined with the topographic differences between movie-watching and resting-state event activity, strongly suggests that co-fluctuation amplitude is at least in part modulated by subjects’ cognitive states.

### FC events enhance identifiability

In the previous sections, we showed that FC events contribute disproportionately to the brain’s static FC patterns, shape its modular structure, are underpinned by a population-level mode of brain activity, and synchronize during viewing of naturalistic stimuli. Do events also enhance the identifiability of individual subjects? That is, does FC estimated from events bear a stronger signature of a subject than FC estimated from non-events?

To test this question, we calculated the co-fluctuation amplitude at each time point (Fig. 4a,i). We then isolated the frames with the highest amplitude and estimated FC using only these frames (Fig. 4a,ii). Repeating this procedure for all subjects and scans resulted in a set of 100 feature vectors (10 subjects × 10 scans) that encode a subjects’ FC patterns (Fig. 4a,iii). We then compute the [100 × 100] similarity matrix - otherwise known as the identifiability matrix - and compare the mean within-subject similarity to the mean between-subject similarity (Fig. 4a,iv). This measure - the differential identifiability - indicates how much more similar subjects’ FC patterns are to themselves than to those of other subjects. We repeat this entire procedure using only low-amplitude frames (Fig. 4a,v-vii), and compare low- and high-amplitude differential identifiability.

**FIG. 4.**
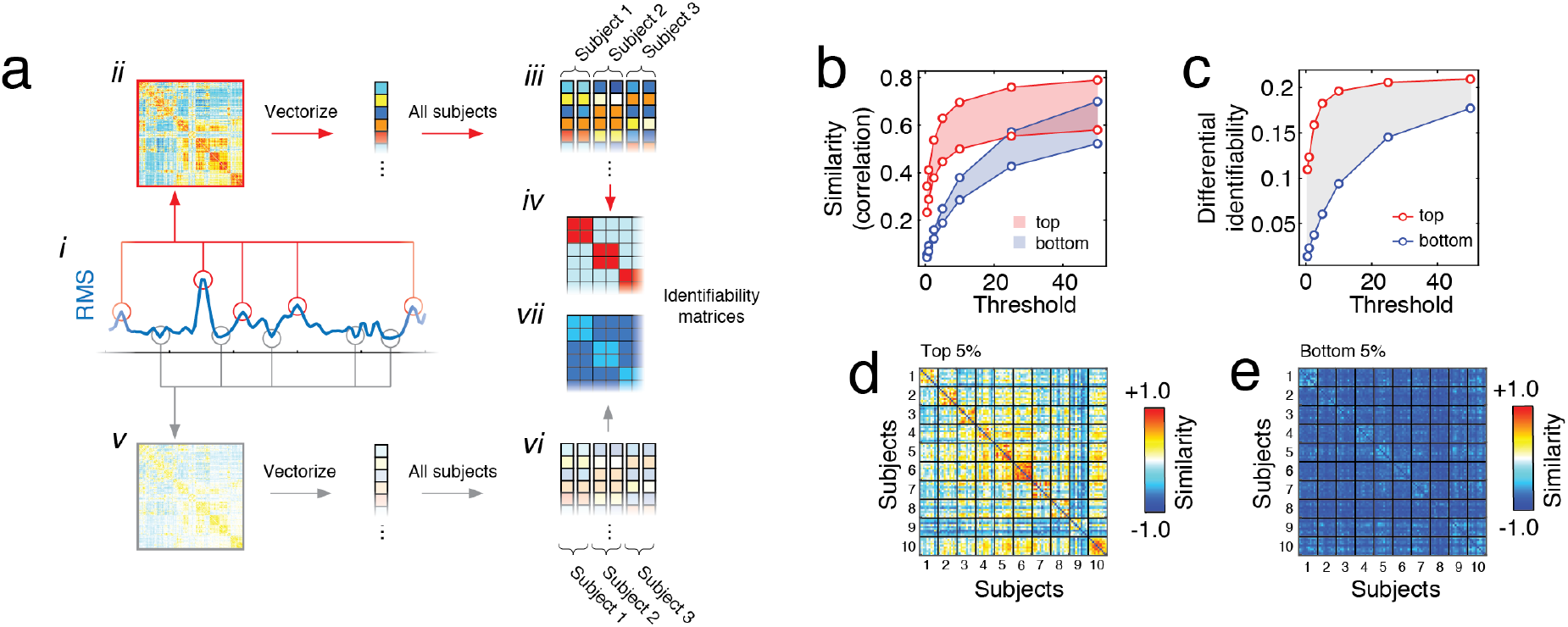
Connectome fingerprints are strong during events and weak during non-events. We investigated whether subject-specific features of rsFC were more prevalent during events or non-events. To address this question, we identified frames (time points) with the highlight and lowest co-fluctuation amplitude and estimated subjects FC using these data, only. We then calculated the inter-subject similarity matrix, i.e. the identifiability matrix. Panel *a* illustrates this general procedure, beginning by isolating high-amplitude time points (*i*), estimation of FC (*ii*), repeating this procedure for all subjects (*iii*), and estimating the inter-subject similarity matrix (*iv*). An identical procedure was carried out for low-amplitude frames and is illustrated in panels *v-vii*. (*b*) We calculated the mean within- and between-subject similarity using both the top (red) and bottom (blue) frames, ordered by co-fluctuation amplitude. For each set of frames, we produce two separate curves, one for the within-subject similarity and another for the between subject similarity. The area between the curves is the *differential identifiability,* or the extent to which subjects’ FC patterns are more similar to themselves than to FC estimated from other subjects. (*c*) We found that differential identifiability was always greater when FC was estimated using the top frames, ordered by amplitude. For the sake of visualization, we show identifiability matrices estimated using high- and low-amplitude frames in panels *d* and *e*, respectively.

In general, we find that inter-subject similarity is greater during events than during non-events and over a range of event thresholds (Fig. 4b). Importantly, we also find that the gap between within- and between-subject similarity, differential identifiability, is greater for events (Fig. 4c-e). This observation suggests that events carry more individualized and distinguishable information about subjects than non-events.

## DISCUSSION

Here, we presented a general approach for “temporally unwrapping” Pearson correlations to generate time series of inter-regional co-fluctuations along network edges. This simple procedure enables us to parse the contributions made by individual frames to rsFC. We find that, in general, we can accurately estimate whole-brain rsFC and its system-level organization using data from a relatively small number of frames. Importantly, we link these frames to a high-modularity brain state and to a specific mode of brain activity, in which default mode and control networks fluctuate in opposition to sensorimotor and attention systems. Our results also suggest that the co-fluctuation patterns, at a coarse scale, capture cognitively relevant fluctuations in brain state and that “events” encode signatures of an individual.

### Decomposing static FC into co-fluctuation snapshots

Central to this paper is the observation that static FC can be non-parametrically decomposed into a series of time-varying “snapshots,” each of which expresses an instantaneous pattern of inter-regional co-fluctuation. Critically, the average of these patterns across time is *exactly* equal to whole-brain static FC. This mathematical truism allows us to neatly assess the contributions of momentary co-fluctuations to the overall FC pattern and to establish a clear link between FC and fluctuations in brain activity.

Our findings, which complement previous work [15, 25– 27], suggest that static FC is driven by contributions from relatively few time points namely those with the highest levels of co-fluctuation amplitude. Frames with low levels of co-fluctuation, on the other hand, contribute little. Because the co-fluctuation time series are estimated at a temporal resolution of single frames, we directly compared network-level events with coincident patterns of brain activity. We established that, at rest, events occur due a specific mode of brain activity that emphasizes oppositional activation of sensorimotor and association cortex.

These observations both clarify and challenge core assumptions concerning static FC and brain network dynamics [28]. Specifically, our findings suggest that wholebrain FC follows a “bursty” trajectory through a highdimensional state space, with extended periods of quietude punctuated by brief and intermittent “events,” whose timing is not clearly related to motion or physiological artifacts. This observation leads to several questions, the most important of which concerns the origins of events. Are events spontaneous occurrences? Are they relevant, in any way, to ongoing cognitive processes? How individualized are events?

### Linking co-fluctuation events to cognition and individual differences

We performed two separate analyses in order to help clarify the origins of events. First, we performed a comparison of event structure at rest and during movie-watching [22]. While the amplitude of co-fluctuations was not statistically different between conditions, we found that co-fluctuations were correlated during movie-watching, suggesting that “events” were being driven by audiovisual features in movies. This finding supports the hypothesis that the timing of events is linked to perception and processing of sensory information, and further suggests that events are not simply spontaneous occurrences. These observation open up possibilities for future studies, that leverage the temporal structure of co-fluctuation amplitudes to track changes in an individual’s cognitive state across time.

Importantly, we also discover differences in the mode of brain activity underpinning events in movie-watching compared to rest. In particular, we found stronger expression of visual and dorsal attention networks, brain systems that one might hypothesize to play an important role in processing visual information and redirecting attentional resources while viewing complex naturalistic stimuli [29, 30]. This finding also demonstrates that the character of events can be modulated by tasks, presenting an opportunity for future studies to comprehensively map the task-evoked topography of events [31].

In our second analysis, we asked whether events were personalized and idiosyncratic [21, 32, 33]. To address this question, we estimated subjects’ FC separately using “event” and “non-event” frames and compared these networks in terms of their *differential identifiability* - the extent to which the similarity of FC patterns was stronger within-subjects than between [34]. Surprisingly, we found that identifiability was significantly stronger during events than non-events, suggesting that subject-specific information is expressed more strongly during event frames.

Collectively, these findings suggest that event structure is highly organized. It tracks time-varying fluctuations in cognitive state and is deeply personalized. These are key observations with clear implications for the study of brain-behavior associations, clinical neuroscience, and phenotype discovery, where the ability to make inferences is limited by the amount of data available. Our results suggest that, by capitalizing on the fact that events carry more subject-specific information than non-events, it may be possible to generate robust network-level biomarkers using a relatively small number of frames and reducing the amount of required data [35] (we note this concept is being explored with other imaging modalities [36]). This approach may be especially useful in clinical and developmental neuroscience, that study populations with characteristics that generally prohibit the extended scan durations necessary for stably estimating FC [37].

### System-level organization emergences from event structure

Lastly, our findings hint at a crucial link between instantaneous fluctuations in activity and the organization of rsFC [38, 39]. Many studies have found that the community structure of rsFC resembles known co-activation patterns, including task-evoked activity [40, 41]. Here, we proposed a strategy that enables us to tease apart the precise contribution of instantaneous BOLD fluctuations (and their topography) to rsFC.

We demonstrated that a particular pattern of activity involving default mode and control regions is primarily responsible for driving co-fluctuation events and, in turn, whole-brain rsFC. While this mode made the greatest contribution, it is likely that other modes make non-trivial contributions as well. By extending the definition of an event to include lower-amplitude fluctuations, we expect to find patterns of activity that correspond to other, well-known brain systems [14]. Moreover, we speculate that these patterns likely recombine in different proportions as a function of task complexity and domain [40, 42] and across individuals [33]. In future work, the proportion of variance explained by different patterns and other statistics related to events, including the frequency with which they occur, may serve as potent correlates of cognitive and disease state. Because events appear to drive the overall configuration of rsFC, we further speculate that their statistics may serve as important complements to traditional measures of rsFC.

### Future work

The approach developed here presents several exciting opportunities for future studies. These include investigating time-varying FC using co-fluctuation patterns, which provide framewise estimates of network structure and circumvent limitations of sliding window approaches [7, 17, 43]. Other possibilities include mapping the relationship of structural connectivity to regional fluctuations or inter-regional co-fluctuations during events and non-events [44] and studying individual differences in cognitive, development, and disease state based on features extracted from events, which we show provide more reliable estimates of subject-level networks.

Importantly, the entire co-fluctuation time series enterprise could be extended in several important ways, including by applying it to other imaging modalities, e.g. electrophysiological recordings [45–48] or fluorescence imaging data [49, 50]. Additionally, it would straightforward to calculate co-fluctuation time series after partialing out the effects of activity elsewhere in the brain [19] or to investigate temporal dependencies and lags between brain regions [51, 52].

### Relationship with existing approaches

We note that the analysis of co-fluctuation time series is conceptually similar to several existing methods [25–27, 53], or in some cases even builds upon shared mathematical machinery [54]. For instance, “multiplication of temporal derivatives” (MTDs) [55] calculates the element-wise products using *differenced* activity time series for all pairs of nodes. These time series are then convolved with a kernel to generate smooth estimates of time-varying nFC. Though similar, our approach relies on untransformed activity to estimate edge time series, thereby preserving the relationship between static nFC and the mean value of each edge time series. Furthermore, our approach omits the smoothing step, making it, in principle, capable of detecting fluctuations in network structure over shorter timescales compared to MTDs.

Another related method is “Co-activation Patterns” (CAPs) [14, 15], which extracts and clusters voxel- or vertex-level activity during high-activity frames. Because a voxel can be co-active under different contexts, the cluster centroids spatially overlap with one another. Though both CAPs and eFC result in overlapping structures, they operate on distinct substrates, with CAPs focusing on activity and eFC focusing on similarity of co-activity. While CAPs requires the specification of additional parameters compared to eFC, e.g. the threshold for a high-activity frame, CAPs may scale better due to the focus on activity rather than connectivity.

Although these approaches arrive at similar conclusions, they possess distinct advantages and disadvantages that make some methods uniquely well-suited for testing specific hypotheses and research questions. For instance, MTDs and the event-based analysis of co-fluctuation time series presented here are appropriate for tracking patterns of connectivity across time. In the case of co-fluctuation time series, which are mathematically related to the static FC pattern, our approach is especially well-suited for assessing the contributions of framewise co-fluctuation patterns to the brain’s overall FC (we note that this relationship, to our knowledge, has not been previously discussed in the extant literature). CAPS and iCAPs, on the other hand, are better-suited for studying activity patterns and tracking their co-occurrences across time. In principle, a systematic and careful comparison of these methods could be carried out in future work.

### Limitations

One of the most key limitations concerns the calculation and interpretation of co-fluctuation time series. The procedure for calculating edge time series begins by z-scoring each brain region’s activity time series. This procedure, however, is only appropriate if the sample mean and standard deviation are temporally invariant [56]. If there is a sustained increase or decrease in activity, e.g. the effect of a blocked task, then the z-scoring procedure can result in a biased mean and standard deviation, resulting in poor estimates of fluctuations in activity. To minimize the likelihood of this occurring, we focused on resting-state and movie-watching data rather than blocked tasks. In future work, investigation of task-evoked co-fluctuations could be investigated by employing already common preprocessing steps, e.g. constructing task regressors to remove the first-order effect of tasks on activity [57].

### Conclusion

In conclusion, our study discloses a novel a link between cortical activity and rsFC, facilitating a statistical explanation of the brain’s system-level architecture in terms of intermittent, short-lived, high-amplitude fluctuations in activity and co-activity. Our methodological framework is readily applicable to other imaging datasets and recording modalities, including observations at neuronal scales, enabling the study of neural co-activity at unprecedented temporal resolution.

## MATERIALS AND METHODS

### Datasets

We analyzed three separate datasets. Specifically, we focused on resting-state data from both The Human Connectome Project and Midnight Scan Club. These data were processed similarly, the details of which are described in this section. The third dataset, which has been analyzed elsewhere [22], includes both resting-state and movie-watching data from a cohort of 29 individuals. This dataset was processed separately using a different procedure and is described in its own section.

The Human Connectome Project (HCP) dataset [20] included resting state functional data (rsfMRI) from 100 unrelated adult subjects (54% female, mean age = 29.11 ± 3.67, age range = 22-36). The study was approved by the Washington University Institutional Review Board and informed consent was obtained from all subjects. Subjects underwent four 15 minute rsfMRI scans over a two day span. A full description of the imaging parameters and image prepocessing can be found in [58]. The rsfMRI data was acquired with a gradient-echo EPI sequence (run duration = 14:33 min, TR = 720 ms, TE = 33.1 ms, flip angle = 52°, 2 mm isotropic voxel resolution, multiband factor = 8) with eyes open and instructions to fixate on a cross. Images were collected on a 3T Siemens Connectome Skyra with a 32-channel head coil.

The Midnight Scan Club (MSC) dataset [21] included rsfMRI from 10 adults (50% female, mean age = 29.1 ± 3.3, age range = 24-34). The study was approved by the Washington University School of Medicine Human Studies Committee and Institutional Review Board and informed consent was obtained from all subjects. Subjects underwent 12 scanning sessions on separate days, each session beginning at midnight. 10 rsfMRI scans per subject were collected with a gradient-echo EPI sequence (run duration = 30 min, TR = 2200 ms, TE = 27 ms, flip angle = 90^°^, 4 mm isotropic voxel resolution) with eyes open and with eye tracking recording to monitor for prolonged eye closure (to assess drowsiness). Images were collected on a 3T Siemens Trio.

### Image Preprocessing of HCP and MSC datasets

#### HCP Functional Preprocessing

Functional images in the HCP dataset were minimally preprocessed according to the description provided in [58]. Briefly, these data were corrected for gradient distortion, susceptibility distortion, and motion, and then aligned to a corresponding T1-weighted (T1w) image with one spline interpolation step. This volume was further corrected for intensity bias and normalized to a mean of 10000. This volume was then projected to the *32k_fs_LR* mesh, excluding outliers, and aligned to a common space using a multi-modal surface registration [59]. The resultant CIFTI file for each HCP subject used in this study followed the file naming pattern: *_REST{1,2}_{L,R}_Atlas_MSMAll.dtseries.nii.

#### MSC Functional Preprocessing

Functional images in the MSC dataset were preprocessed using *fMRIPrep* 1.3.2 [60], which is based on Nipype 1.1.9 [61]. The following description of *fMRIPrep*’s preprocessing is based on boilerplate distributed with the software covered by a “no rights reserved” (CC0) license. Internal operations of *fMRIPrep* use Nilearn 0.5.0 [62], ANTs 2.2.0, FreeSurfer 6.0.1, FSL 5.0.9, and AFNI v16.2.07. For more details about the pipeline, see the section corresponding to workflows in *fMRIPrep* ‘s documentation.

The T1-weighted (T1w) image was corrected for intensity non-uniformity with **N4BiasFieldCorrection** [63, 64], distributed with ANTs, and used as T1w-reference throughout the workflow. The T1w-reference was then skull-stripped with a Nipype implementation of the **antsBrainExtraction**.sh workflow, using NKI as the target template. Brain surfaces were reconstructed using recon-all [65], and the brain mask estimated previously was refined with a custom variation of the method to reconcile ANTs-derived and FreeSurfer-derived segmentations of the cortical gray-matter using Mindboggle [66]. Spatial normalization to the *ICBM 152 Nonlinear Asymmetrical template version 2009c* [67] was performed through nonlinear registration with antsRegistration, using brain-extracted versions of both T1w volume and template. Brain tissue segmentation of cerebrospinal fluid (CSF), white-matter (WM) and gray-matter (GM) was performed on the brain-extracted T1w using FSL’s fast [68].

Functional data was slice time corrected using AFNI’s 3dTshift and motion corrected using FSL’s mcflirt [69]. *Fieldmap-less* distortion correction was performed by co-registering the functional image to the same-subject T1w image with intensity inverted [70] constrained with an average fieldmap template [71], implemented with antsRegistration. This was followed by co-registration to the corresponding T1w using boundary-based registration [72] with 9 degrees of freedom. Motion correcting transformations, field distortion correcting warp, BOLD-to-T1w transformation and T1w-to-template (MNI) warp were concatenated and applied in a single step using **antsApplyTransforms** using Lanczos interpolation.

Several confounding time-series were calculated based on this preprocessed BOLD: framewise displacement (FD), DVARS and three region-wise global signals. FD and DVARS are calculated for each functional run, both using their implementations in Nipype [73]. The three global signals are extracted within the CSF, the WM, and the whole-brain masks.

The resultant NIFTI file for each MSC subject used in this study followed the file naming pattern **\*_space-T1w_desc-preproc_bold.nii.gz**.

### Image Quality Control

All functional images in the HCP dataset were retained. The quality of functional images in the MSC were assessed using *fMRIPrep*’s visual reports and *MRIQC* 0.15.1 [74]. Data was visually inspected for whole brain field of view coverage, signal artifacts, and proper alignment to the corresponding anatomical image. Functional data were excluded if greater than 25% of the frames exceeded 0.2 mm framewise displacement [75]. Furthermore, functional data were excluded if marked as an outlier (exceeding 1.5x inter-quartile range in the adverse direction) in more than half of the following image quality metrics (calculated within-dataset, across all functional acquisitions): *dvars, tsnr, fd_mean, aor, aqi, snr,* and *efc.* Information about these image quality metrics can be found within *MRIQC*’s documentation [76].

### Functional and Structural Networks Preprocessing

#### Parcellation Preprocessing

A functional parcellation designed to optimize both local gradient and global similarity measures of the fMRI signal [77] *(Schaefer200*) was used to define 200 areas on the cerebral cortex. These nodes are also mapped to the *Yeo* canonical functional networks [38]. For the HCP dataset, the *Schaefer200* is openly available in *32k_fs_LR* space as a CIFTI file. For the MSC and HBM datasets, a *Schaefer200* parcellation was obtained for each subject using a Gaussian classifier surface atlas [78] (trained on 100 unrelated HCP subjects) and FreeSurfer’s mris_ca_label function. These tools utilize the surface registrations computed in the recon-all pipeline to transfer a group average atlas to subject space based on individual surface curvature and sulcal patterns. This method rendered a T1w space volume for each subject. For use with functional data, the parcellation was resampled to 2mm T1w space.

#### Functional Network Preprocessing

The mean BOLD signal for each cortical node data was linearly detrended, band-pass filtered (0.008-0.08 Hz) [75], confound regressed and standardized using Nilearn’s signal.clean, which removes confounds orthogonally to the temporal filters [79]. The confound regression employed [80] included 6 motion estimates, time series of the mean CSF, mean WM, and mean global signal, the derivatives of these nine regressors, and the squares these 18 terms. Furthermore, a spike regressor was added for each fMRI frame exceeding a motion threshold (HCP = 0.25 mm root mean squared displacement, MSC = 0.5 mm framewise displacement). This confound strategy has been shown to be relatively effective option for reducing motion-related artifacts [75]. Following this preprocessing and nuisance regression, residual mean BOLD time series at each node was recovered.

### Image Preprocessing of Indiana University Dataset

#### Demographics

We analyzed MRI data collected from *N_s_* = 29 subjects (5 female, 24 male; 25 were right-handed). This cohort was male-dominant, as subjects were intended to serve as controls for a study in autism spectrum disorder, which is more common in men than women. At the time of their first scan, the average subject age was 24.9 ± 4.7 years [81].

#### MRI acquisition and processing

MRI images were acquired using a 3T whole-body MRI system (Magnetom Tim Trio, Siemens Medical Solutions, Natick, MA) with a 32-channel head receive array. Both raw and prescan-normalized images were acquired; raw images were used at all preprocessing stages and in all analyses unless specifically noted. During functional scans, T2*-weighted multiband echo planar imaging (EPI) data were acquired using the following parameters: TR/TE = 813/28 ms; 1200 vol; flip angle = 60°; 3.4 mm isotropic voxels; 42 slices acquired with interleaved order covering the whole brain; multi-band acceleration factor of 3. Preceding the first functional scan, gradient-echo EPI images were acquired in opposite phase-encoding directions (10 images each with P-A and A-P phase encoding) with identical geometry to the EPI data (TR/TE = 1175/39.2 ms, flip angle = 60°) to be used to generate a fieldmap to correct EPI distortions, similar to the approach used by the Human Connectome Project [82]. High-resolution T1-weighted images of the whole brain (MPRAGE, 0.7 mm isotropic voxel size; TR/TE/TI = 2499/2.3/1000 ms) were acquired as anatomical references.

All functional data were processed according to an in-house pipeline using FEAT (v6.00) and MELODIC (v3.14) within FSL (v.5.0.9; FMRIB’s Software Library, www.fmrib.ox.ac.uk/fsl), Advanced Normalization Tools (ANTs; v2.1.0) [83], and Matlab_R2014b. This pipeline was identical to the **GLM + MGTR** procedure described in [84].

In more detail, individual anatomical images were bias-corrected and skull-stripped using ANTs, and segmented into gray matter, white matter, and CSF partial volume estimates using FSL FAST. A midspace template was constructed using ANTs’ *buildtemplateparallel* and subsequently skull-stripped. Composite (affine and diffeomorphic) transforms warping each individual anatomical image to this midspace template, and warping the midspace template to the Montreal Neurological Institute MNI152 1mm reference template, were obtained using ANTs.

For each functional run, the first five volumes (≈4 seconds) were discarded to minimize magnetization equilibration effects. Framewise displacement traces for this raw (trimmed) data were computed using *fsLmotion_outliers.* Following [84, 85], we performed FIX followed by mean cortical signal regression. This procedure included rigid-body motion correction, fieldmap-based geometric distortion correction, and non-brain removal (but not slice-timing correction due to fast TR [82]). Preprocessing included weak highpass temporal filtering (>2000 s FWHM) to remove slow drifts [82] and no spatial smoothing. Off-resonance geometric distortions in EPI data were corrected using a fieldmap derived from two gradient-echo EPI images collected in opposite phase-encoding directions (posterior-anterior and anterior-posterior) using FSL topup.

We then used FSL-FIX [86] to regress out independent components classified as noise using a classifier trained on independent but similar data and validated on hand-classified functional runs. The residuals were regarded as “cleaned” data. Finally, we regressed out the mean cortical signal (mean BOLD signal across gray matter partial volume estimate obtained from FSL FAST). All analyses were carried out on these data, which were registered to subjects’ skull-stripped T1-weighted anatomical imaging using Boundary-Based Registration (BBR) with *epi_reg* within FSL. Subjects’ functional images were then transformed to the MNI152 reference in a single step, using ANTS to apply a concatenation of the affine transformation matrix with the composite (affine + diffeomorphic) transforms between a subject’s anatomical image, the midspace template, and the MNI152 reference. Prior to network analysis, we extracted mean regional time series from regions of interest defined as sub-divisions of the 17-system parcellation reported in [38] and used previously [4, 87, 88]. Wakefulness during movie and rest scans was monitored in real-time using an eye tracking camera (Eyelink 1000).

#### Naturalistic stimuli

All movies were obtained from Vimeo (https://vimeo.com). They were selected based on multiple criteria. First, to ensure that movies represented novel stimuli, we excluded any movie that had a wide theatrical release. Secondly, we excluded movies with potentially objectionable content including nudity, swearing, drug use, etc. Lastly, we excluded movies with intentionally startling events that could lead to excessive in-scanner movement.

Each movie lasted approximately 1 to 5 minutes. Each movie scan comprised between four and six movies with genres that included documentaries, dramas, comedies, sports, mystery, and adventure. See Table. S1 for more details.

### Co-fluctuation time series

Constructing networks from fMRI data (or any neural time series data) requires estimating the statistical dependency between every pair of time series. The magnitude of that dependency is usually interpreted as a measure of how strongly (or weakly) those voxels are parcels are functionally connected to each other. By far the most common measure of statistic dependence is the Pearson correlation coefficient. Let x*_i_* = [x*_i_*(1),…, x*_i_*(*T*)] and X*_j_* = [x*_j_*(1),…, x*_j_*(*T*)] be the time series recorded from voxels or parcels *i* and *j*, respectively. We can calculate the correlation of *i* and *j* by first z-scoring each time series, such that 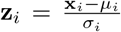, where 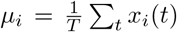 and 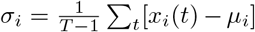 are the time-averaged mean and standard deviation. Then, the correlation of *i* with *j* can be calculated as: 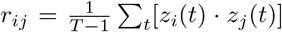. Repeating this procedure for all pairs of parcels results in a node-by-node correlation matrix, i.e. an estimate of FC. If there are *N* nodes, this matrix has dimensions [*N × N*].

To estimate *edge*-centric networks, we need to modify the above approach in one small but crucial way. Suppose we have two z-scored parcel time series, **z***_i_* and **Z***_j_*. To estimate their correlation we calculate the mean their element-wise product (not exactly the average, because we divide by *T* − 1 rather than *T*). Suppose, instead, that we never calculate the mean and simply stop after calculating the element-wise product. This operation would result in a vector of length T whose elements encode the moment-by-moment co-fluctuations magnitude of parcels *i* and *j.* For instance, suppose at time t, parcels *i* and *j* simultaneously increased their activity relative to baseline. These increases are encoded in **z***_i_* and **Z***_j_* as positive entries in the tth position, so their product is also positive. The same would be true if *i* and *j decreased* their activity simultaneously (because the product of negatives is a positive). On the other hand, if *i* increased while *j* decreased (or *vice versa*), this would manifest as a negative entry. Similarly, if either i or j increased or decreased while the activity of the other was close to baseline, the corresponding entry would be close to zero.

Accordingly, the vector resulting from the element-wise product of **z***_i_* and **Z***_j_* can be viewed as encoding the magnitude of moment-to-moment co-fluctuations between *i* and *j*. An analogous vector can easily be calculated for every pair of parcels (network nodes), resulting in a set of co-fluctuation (edge) time series. With N parcels, this results in 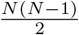 pairs, each of length T.

### Modularity maximization

Modularity maximization is a heuristic for detecting communities in networks [89]. Intuitively, it attempts to decompose a network into non-overlapping sub-networks such that the observed density of connections within sub-networks maximally exceeds what would be expected by chance, where chance is determined by the user. The actual process of detecting communities is accomplished by choosing community assignments that maximize a modularity quality function, *Q,* defined as:

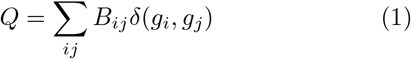

where B*_i,j_* = *A_i,j_* — P*_i,j_* is the *{i,j}* element of the modularity matrix, which represents the observed weight of the connection between nodes *i* and *j* minus the expected weight. The variable g¿ is the community assignment of node *i* and δ(*x, y*) is the Kronecker delta function, whose value is 1 when *g_i_* = *g_j_* and 0 otherwise. The modularity, Q, is effectively a sum over all edges that fall within communities and is optimized when the the observed weights of connections is maximally greater than the expected. In general, larger values of Q are thought to reflect superior community partitions.

#### Signed and correlation matrices

In this manuscript, we used the following variant of modularity, *q\**, which has been shown to be especially well-suited for use with correlation matrices [23]:

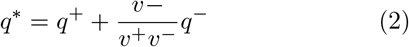

where 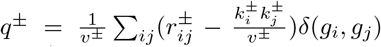. In this expression, 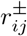 represents either the positive or negative elements of the correlation matrix, 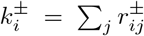, and 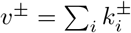.

### Differential identifiability

Let *A_1_* be a *N × N* FC matrix. We can represent this matrix as a *M* = *N* × (N — 1)/2-dimensional vector by extracting its upper triangle elements. We can assess the similarity of two matrices, *A_1_* and *A_2_*, by computing the similarity of their vector representations. Suppose we had multiple scans from multiple individuals. Let *I_self_* and *I_others_* be the average within- and between-subject similarity. Differential identifiability, then, is simply: *I_diff_* = *I_self_* — *I_diff_* [34]. Intuitively, the larger the value of *I_diff_*, the stronger the population-level “fingerprint”

[32].

## AUTHOR CONTRIBUTIONS

RFB, JF, and OS conceived of study. JF and LB processed data. FZE, YJ, JF, LB, DPK, OS, and RB carried out all analyses, wrote, edited, and revised the submitted manuscript.

## ACKNOWLEDGMENTS

RFB and FZE acknowledge support from Indiana University Office of the Vice President for Research Emerging Area of Research Initiative, Learning: Brains, Machines and Children. This work was supported by the NIH (R01MH110630 and R00MH094409 to DPK and T32HD007475 Postdoctoral Traineeship to LB).

## DATA AVAILABILITY

All imaging data come from publicly-available, openaccess repositories. Human connectome project data can be accessed *via* https://db.humanconnectome.org/app/template/Login.vm after signing a data use agreement. Midnight scan club data can be accessed *via* OpenfMRI at https://www.openfmri.org/dataset/ds000224/. The Indiana University dataset is available upon reasonable request.

## CODE AVAILABILITY

All processing and analysis code is available upon reasonable request.

---

**FIG. S1.**
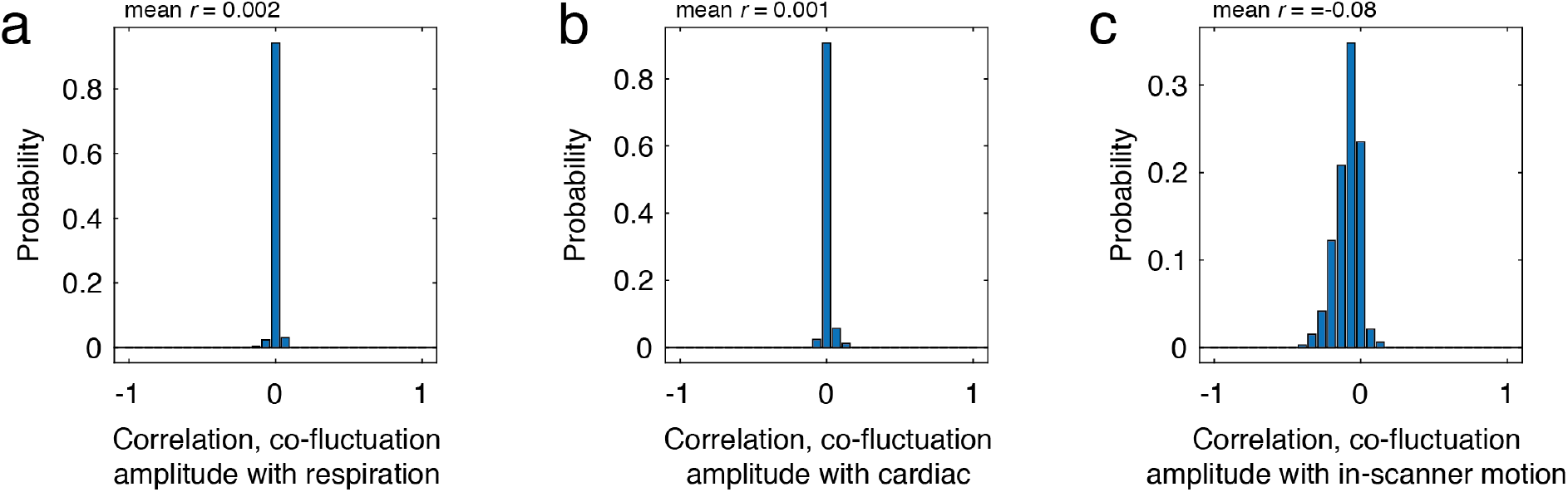
Comparison of co-fluctuation amplitude with confounding variables. In the main text we calculated the magnitude of co-fluctuation at every frame. A concern is that variation in this measure could be attributed to physiological and motion-related variables of non-neural origins. To address this concern, we calculated the correlation of co-fluctuation amplitude with three variables: respiration and cardiac data as well as in-scanner head motion (relative root mean square error framewise displacement). In-scanner motion is already sampled at the same frequency as the BOLD acquisition; for the two physiological variables, we calculated the mean value within a frame. We calculated these variables for every subject and scan session in the HCP dataset and computed their correlation with the co-fluctuation amplitude. The distributions of correlation coefficients, shown here in panels *a*, *b*, and *c*, were tightly centered on zero, suggesting that co-fluctuation amplitude is not obviously related to standard physiological or motion-related variables.

**FIG. S2.**
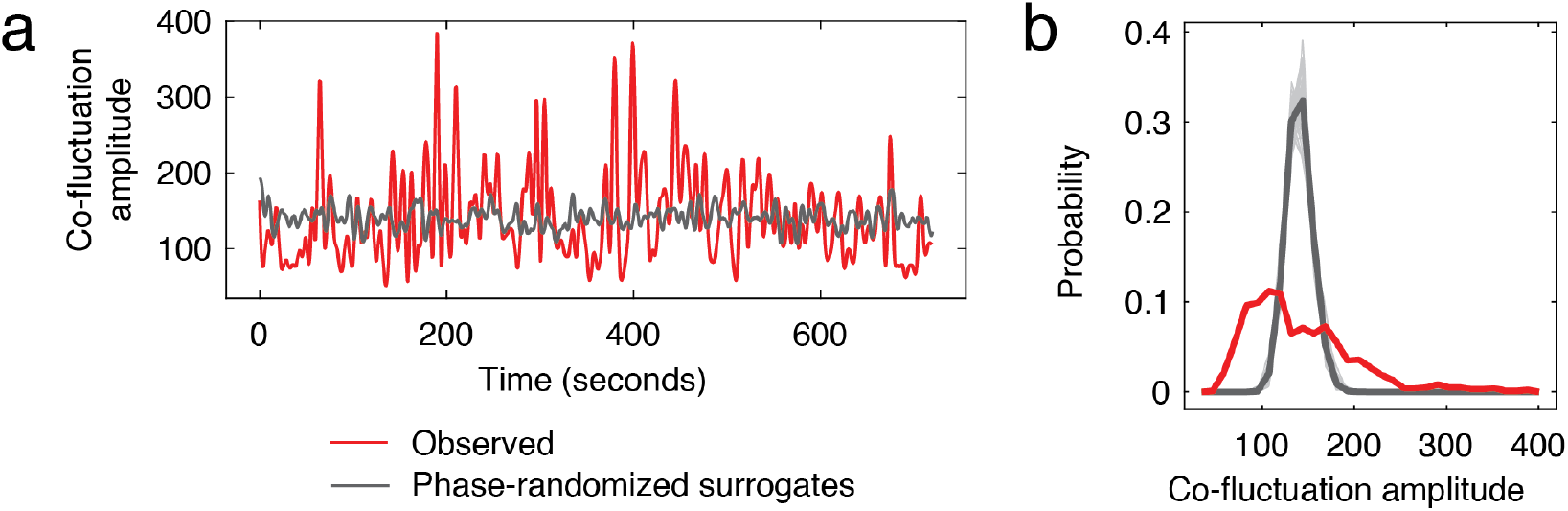
Comparison of co-fluctuation amplitude from observed data and from phase-randomized surrogates. In the main text we calculate co-fluctuation amplitude as root sum square of edge co-fluctuations at each moment in time. Here, we compare these observed amplitudes with those estimated from phase-randomized surrogate time series. The phase randomization procedure has been described in detail elsewhere [90]. Briefly, this procedure entails taking the discrete Fourier transform of each regional BOLD time series, adding random phase at each frequency bin, and taking the inverse Fourier transform, generating a surrogate time series for that region with same power spectrum but random phase properties. We repeat this procedure for all *N* = 200 regions and subsequently calculate their co-fluctuation time series and overall amplitude (panel *a*). We repeated this procedure 100 times and found that the distribution of co-fluctuation amplitude for the observed data was broad and included a heavy tail that was not present in the surrogate data (panel *b*). This observation suggests that the observed co-fluctuations (in particular the high-amplitude “events”) cannot be explained by spectral properties of the fMRI BOLD time series alone.

**FIG. S3.**
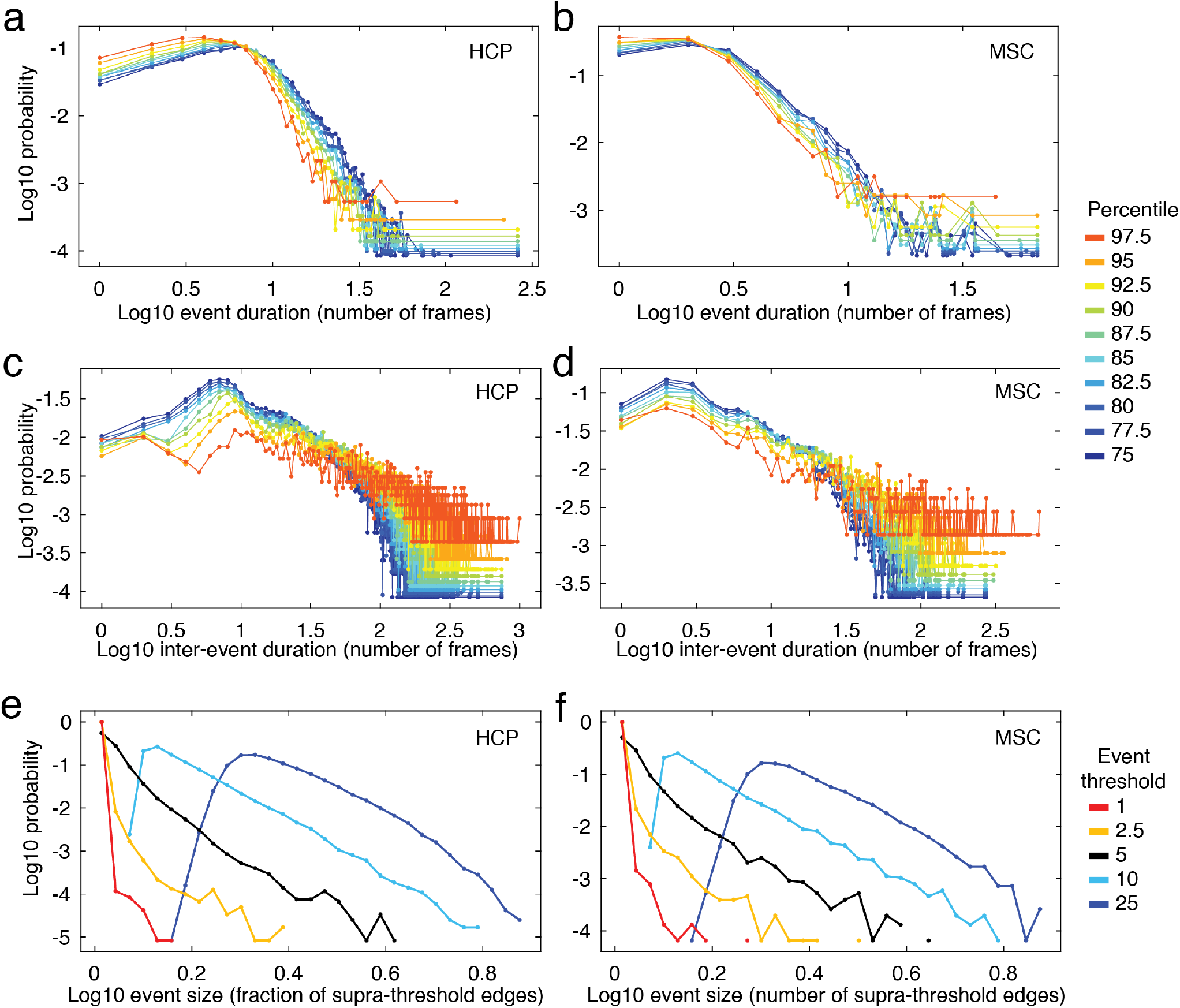
Event duration and inter-event interval distributions. For each scan session, we calculated the co-fluctuation amplitude at every frame. We imposed percentile-based thresholds on these data (percentiles calculated based on pooled data from all subjects and all scan sessions). Thresholding the co-fluctuation amplitude time series results in a binary classification of time points as either “events” or “non-events”. From these observations, we calculated two quantities: “event duration” as the number of consecutive frames classified as events and “inter-event duration” as the number of frames between successive events. We repeated this analysis for both the HCP and MSC datasets. Panels *a* and *b* show event durations for HCP and MSC datasets, respectively. Note that the distribution is broad and includes a heavy tail, indicating a lack of periodicity. Panels *c* and *d* depict inter-event durations for HCP and MSC datasets. Additionally, we assessed the size of events, as measured by the fraction of all edges whose co-fluctuation amplitude at a given frame exceeded some threshold. Here, we identified events as time points at which the co-fluctuation amplitude was in the top 1%, 2.5%, 5%, 10%, and 25% (event thresholds are indicated by different colors in each plot). Then, for each time point classified as an event, we calculated the fraction of all edges whose absolute co-fluctuation amplitude exceed the 75th percentile. We performed this procedure using both HCP (*e*) and MSC *(f*) data and found that event sizes follow a broad and heavy-tailed distribution, suggesting that they follow no characteristic scale of description.

**FIG. S4.**
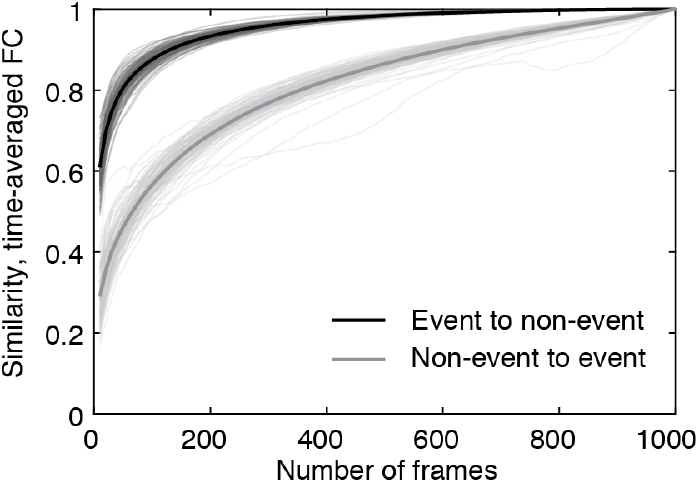
Similarity of time-averaged FC with FC estimated using fewer frames. In the main text, we showed that rsFC, when estimated using the top 5% of frames (ordered by co-fluctuation magnitude) resulted in a connectivity matrix that much more similar to time-averaged FC than the matrix generated using the bottom 5% of frames. Here, we show that this relationship persists irrespective of percentile. To do this for a given subject and scan session, we ordered frames according to co-fluctuation magnitude from greatest to least. Then, we extracted the top and bottom *k* frames (varying *k* from 3 to to *T*, where *T* is the total number of frames in the scan session), estimating FC using those *k* frames, and calculating the similarity with time-averaged FC. This procedure results in a similarity value at every *k* for both the top and bottom frames. We repeated this analysis for all 100 subjects in the HCP dataset. We find that across the full range of *k*, FC estimated using frames corresponding to high-amplitude co-fluctuations was always more similar to the time-averaged FC than those estimated using low-amplitude frames.

**FIG. S5.**
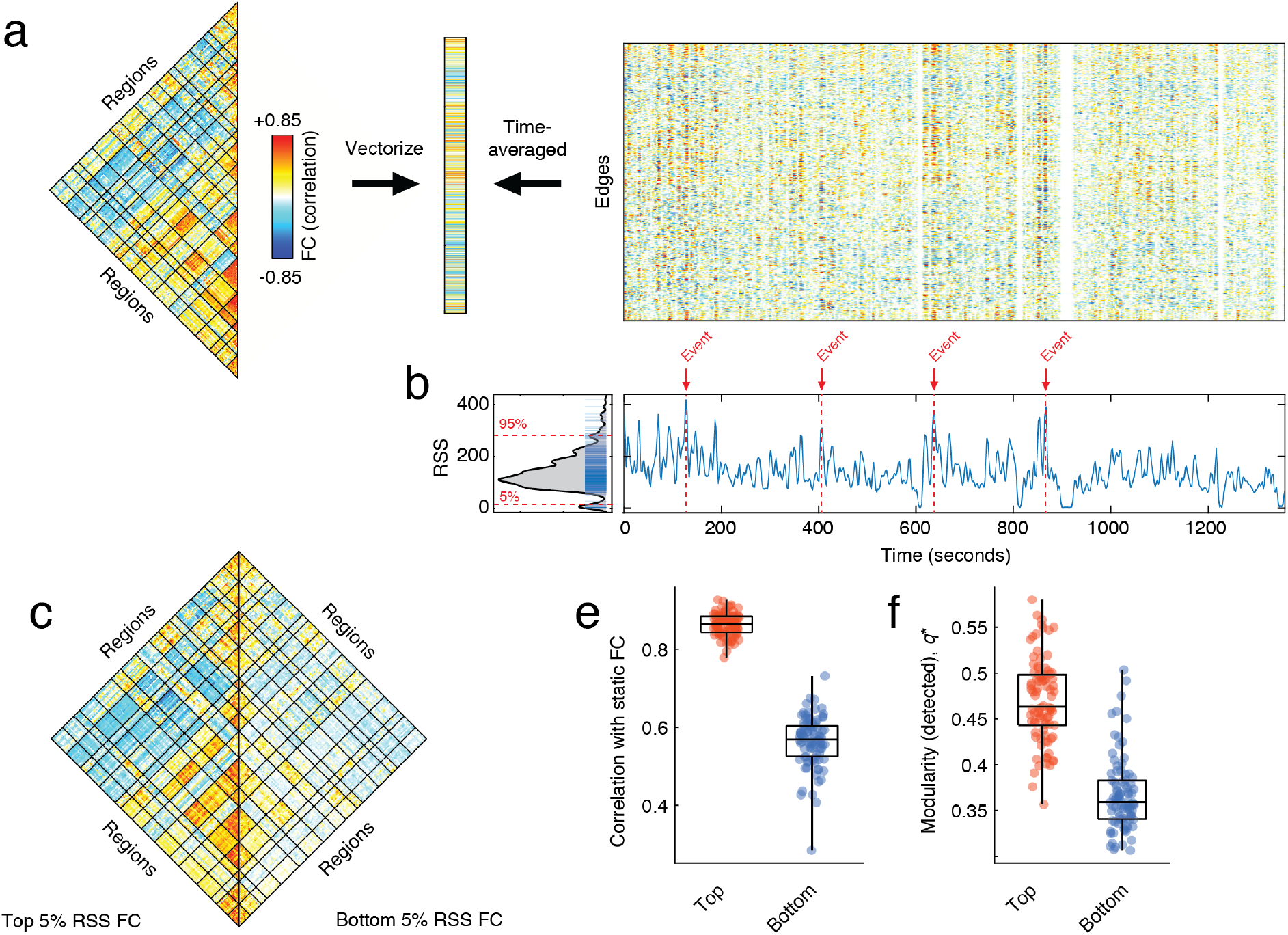
Co-fluctuation time series reveal event structure of resting-state functional connectivity for MSC dataset. (*a*) We use a “temporal unwrapping” of the Pearson correlation to generate co-fluctuation time series for every pair of brain regions (edges). The elements of the co-fluctuation time series are the element-wise products of z-scored regional BOLD time series that, when averaged across time, yield vectors that are exactly equal to the Pearson correlation coefficient and can be rearranged to create a resting-state functional connectivity matrix. (*b*) We find that the co-fluctuation time series contains moments in time where many edges collectively co-fluctuate. We can identify these moments by calculating the root sum square across all co-fluctuation time series and plotting this value as a function of time. We consider high-amplitude values as potential “events”. The distribution of edge co-fluctuation amplitude is heavy tailed. We wanted to assess the contribution of events and non-events to the overall pattern of functional connectivity. To do this, we extracted the top and bottom 5% of all time points (ordered by co-fluctuation amplitude) and estimated functional connectivity from those points alone. (*c*) Average functional connectivity across 100 subjects using top 5% (left) and bottom 5% (right). (*d*) In general, the networks estimated using the top 5% of time points were much more similar to traditional functional connectivity than those estimated using the bottom 5% of time points. (*e*) We performed a similar comparison of network modularity using networks reconstructed using top and bottom 5% frames.

**FIG. S6.**
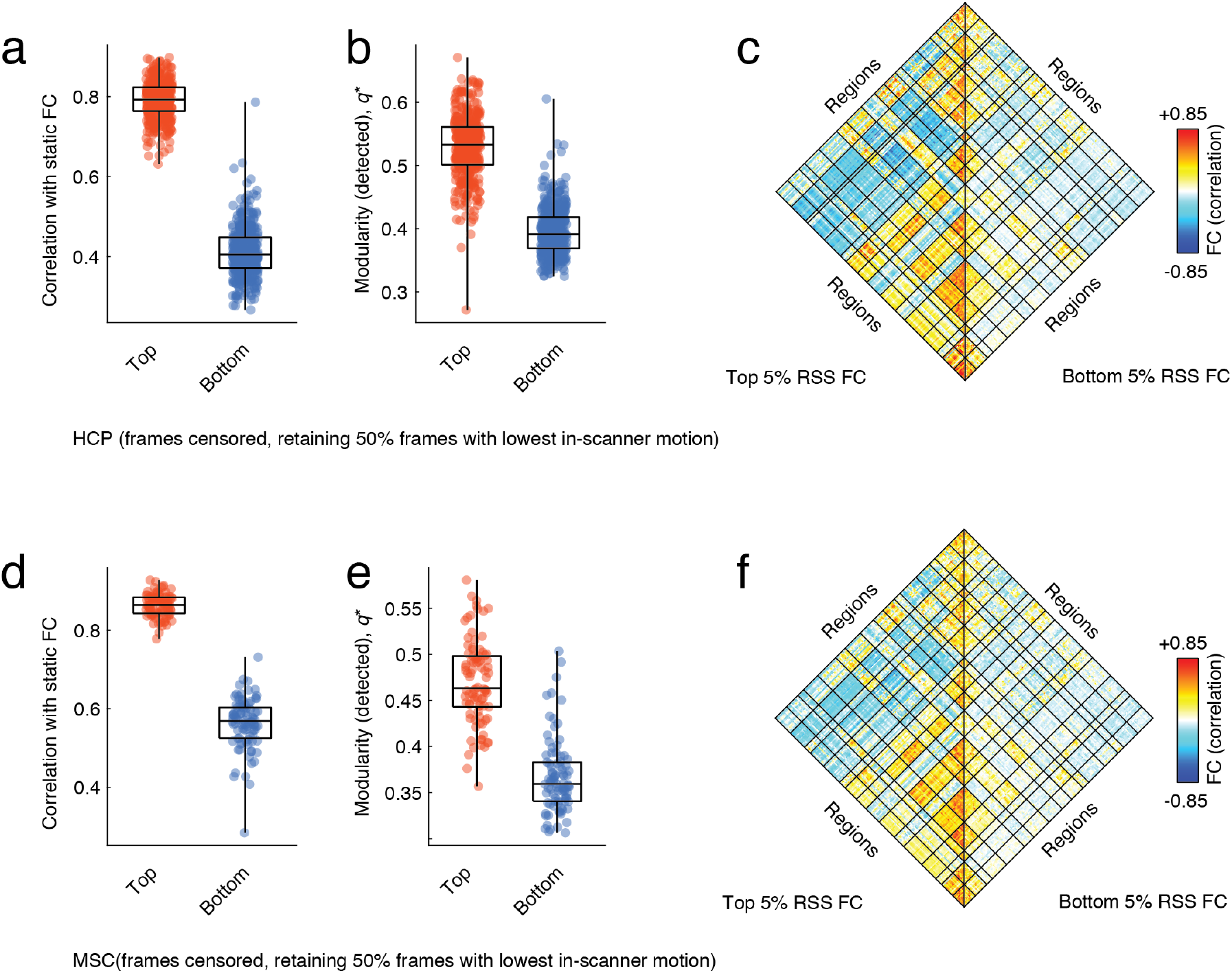
Effect of frame-censoring. In the main text, we demonstrated that FC estimated using frames corresponding to high-amplitude co-fluctuations was more similar to time-averaged FC than the FC estimated using low-amplitude co-fluctuation frames. Here, we perform an identical analysis using only the bottom 50% frames in terms of in-scanner motion [91]. Note that this procedure results in time series that include exactly half of the original frames. (*a*) Correlation of FC estimated using top and bottom 5% of frames, ordered by co-fluctuation amplitude. As in the main text, the top 5% are more correlated with time-averaged FC than the bottom 5%. (*b*) Modularity of FC estimated using only the top and bottom 5% of frames. As in the main text, the top 5% are more modular than the bottom. (*c*) Group-averaged FC matrices estimated using the top 5% of frames (left) and the bottom 5% of frames (right). Panels *a*, *b*, and *c* depict results using HCP data, while *d, e,* and *f* depict analogous results using data from the MSC.

**FIG. S7.**
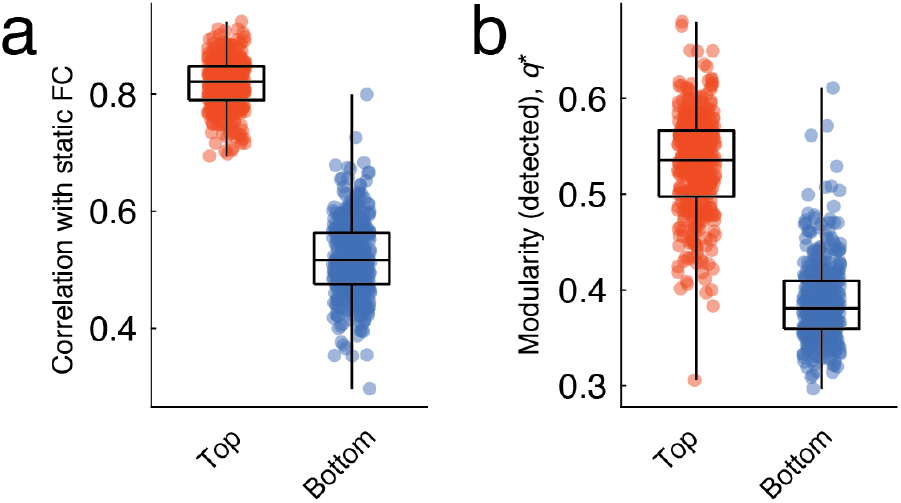
Alternative strategy for estimating FC from a limited number of frames. In the main text, we estimated FC from the top and bottom 5% of frames by extracting fMRI BOLD activity from computing the correlation structure. An alternative strategy for estimating FC is to simply average co-fluctuation time series over the top/bottom frames, ordered by co-fluctuation amplitude. Here, we perform this analysis on HCP data and show that (*a*) FC from the top 5% of frames in terms of co-fluctuation amplitude is more similar to time-averaged FC than FC from the bottom 5% of frames and that (*b*) FC from the top 5% results in more modular networks than FC from the bottom 5%. These results are in exact agreement with what was reported in the main text.

**FIG. S8.**
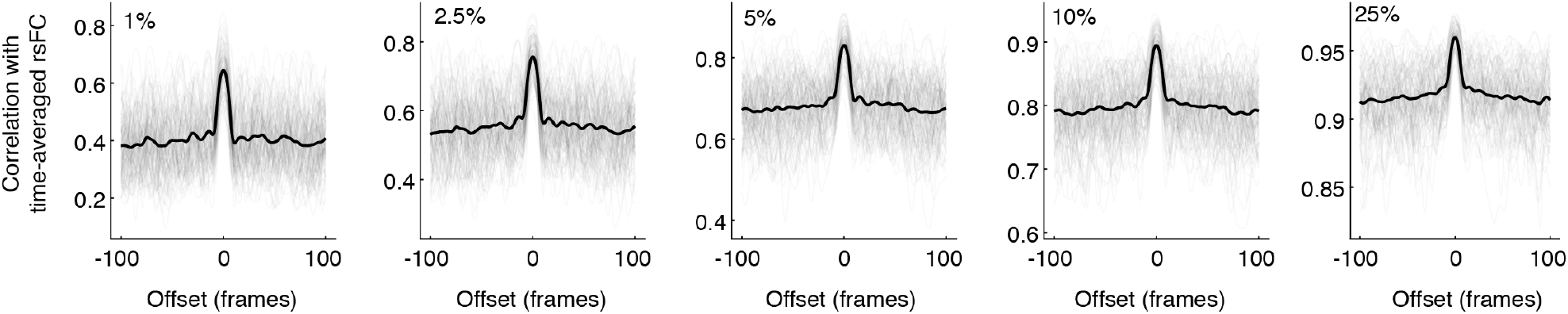
Effect of “jittering” on correspondence between time-averaged rsFC and rsFC estimated using reduced number of frames. In the main text, we demonstrated that the correspondence between time-averaged rsFC and rsFC estimated using high-amplitude co-fluctuation frames was significantly greater than the correspondence using low-amplitude frames. This comparison of the highest- and lowest-amplitude frames can be viewed as a comparison of extremes. A more general test would be to compare the correspondence of rsFC from high-amplitude frames with rsFC from randomly-sampled frames. A truly random sample, however, may destroy any temporal autocorrelation in the time series data. Instead, we identified “events” as frames whose co-fluctuation amplitude exceeded some label, and used the circular shift operator to move these frames forward and backward in time, approximately preserving their temporal structure. Here, we show the correlation of time-averaged rsFC with rsFC estimated using the offset event frames (100 frames forward and backward in time). We repeat this analysis with different event thresholds (from left to right,the top 1%, 2.5%, 5%, 10%, and 25%). In general, we find that the correlation with time-averaged rsFC is peaked exactly at an offset of 0 and rapidly decays to a baseline level. This observation holds for all event thresholds, and suggests that random samples with temporal structure that preserves autocorrelative properties of the co-fluctuation amplitude time series will, in general, result in estimates of rsFC with poorer correspondence to time-averaged rsFC than the event frames, themselves.

**FIG. S9.**
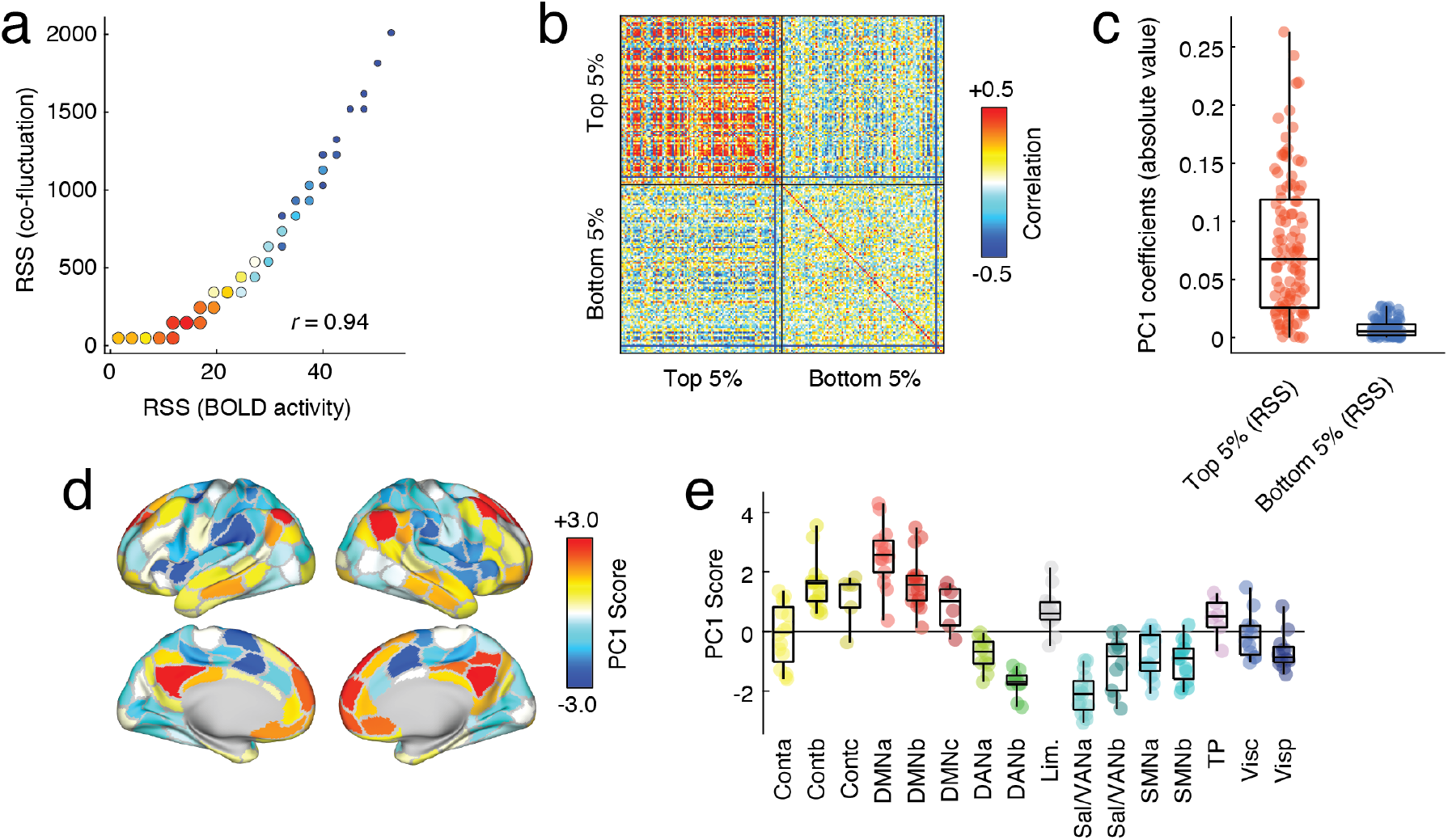
Relationship of network co-fluctuations with BOLD fluctuations for MSC dataset. Here, we replicate results from the main text using the MSC data. Specifically, we relate high-amplitude co-fluctuations to fluctuations in fMRI BOLD activity. We subsequently demonstrate that the high-amplitude fluctuations are driven by activity patterns involving control and default mode networks, and that these patterns are expressed similarly across individuals. As in the main text, we first calculate the root sum square amplitude of BOLD activity at each time point and compare that to the amplitude of co-fluctuations. (*a*) Pooling data from across subjects, we find that these two variables are highly correlated. (*b*) To investigate this relationship further, we extract mean activity patterns for each subject and for each scan during the top and bottom 5% time points, indexed according to co-fluctuation amplitude. Here, we show the correlation matrix of those activity vectors. (*c*) We then performed a principal component analysis of this correlation matrix and found that absolute value of coefficients for the first component (PC1) were greater for the top 5% than the bottom 5%, and (*d*, *e*) the PC1 score corresponded to activity patterns that emphasized correlated fluctuations of default mode and control networks that were weakly or anti-correlated with fluctuations elsewhere in the brain. These observations suggest that co-fluctuation events, which drive resting-state functional connectivity, are underpinned by instantaneous activation and deactivation of default mode and control network areas.

**FIG. S10.**
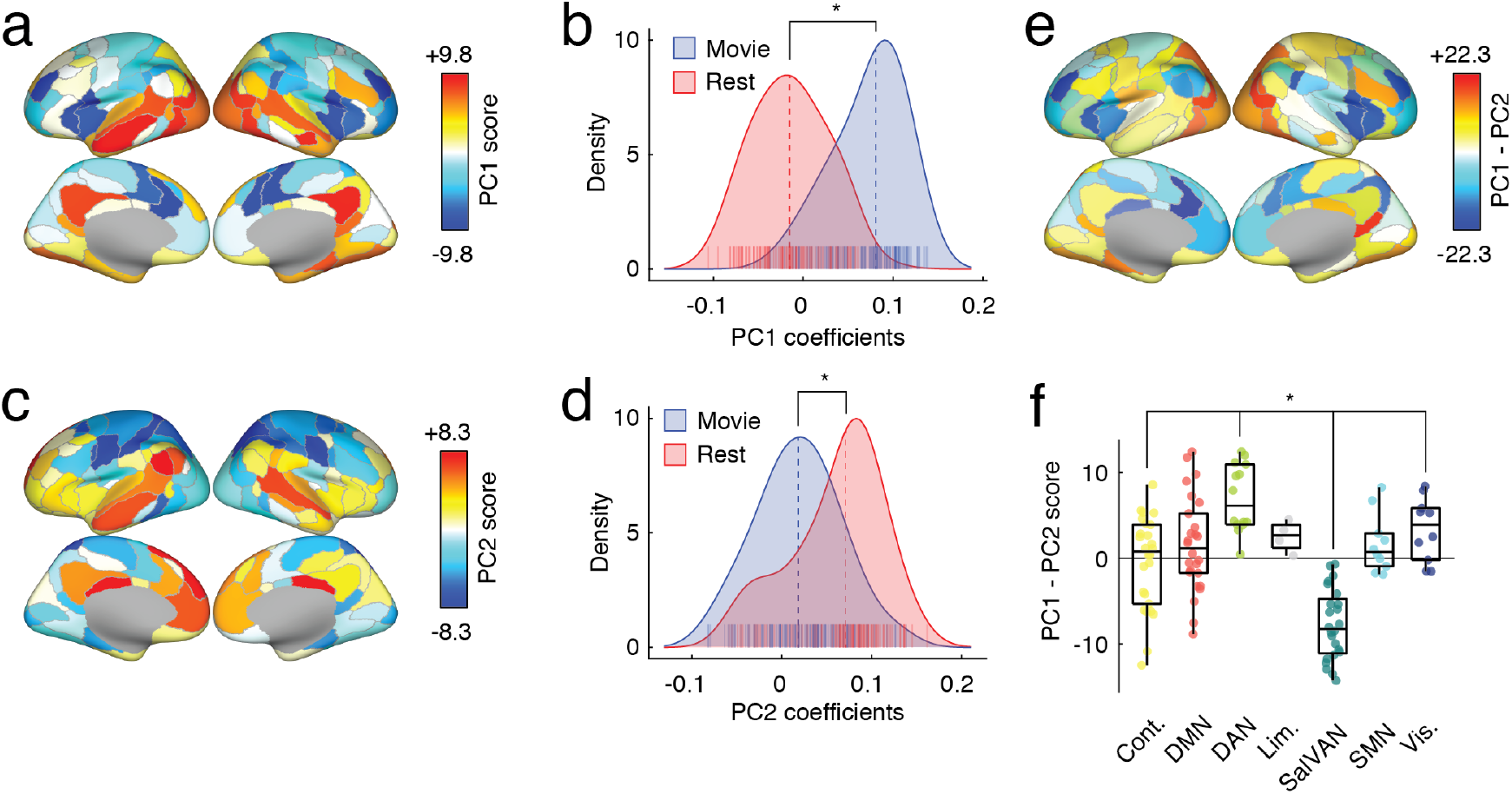
Joint PCA analysis of movie-watching and resting-state data. In the main text we described a procedure for comparing modes of brain activity during events while subjects were either at rest or watching movies. Here, we present an alternative analysis strategy. The procedure in the main text involved identifying activity patterns during high- and low-amplitude frames, concatenating these patterns across subjects, and performing a PCA on the resulting matrix. Importantly, this procedure was carried out separately for resting-state and movie scans. We then compared the first principal component for each condition. Here, we extract high-amplitude frames from resting-state and movie-watching scans, concatenate them in the same matrix, and perform a joint decomposition using PCA. Our analysis focuses on the first two PCs (PC1 and PC2), whose topographic distributions are shown in panels *a* and *c*. We note that the pattern of PC1 is highly similar to the component obtained from our analysis of the movie-watching data described in the main text. Similarly, PC2 is similar to the component obtained from our analysis of the resting-state data described in the main text. Interestingly, we find that the coefficients loading onto PC1 are stronger for movie-watching scans than for resting-state scans (panel *b*), while the opposite is true for PC2 (panel *d*). We highlight regional and system-level differences in panels *e* and *f*. We note that, as in the main text, we find stronger engagement of dorsal attention and visual networks during movie-watching and stronger engagement of control and salience/ventral attention networks at rest.

**FIG. S11.**
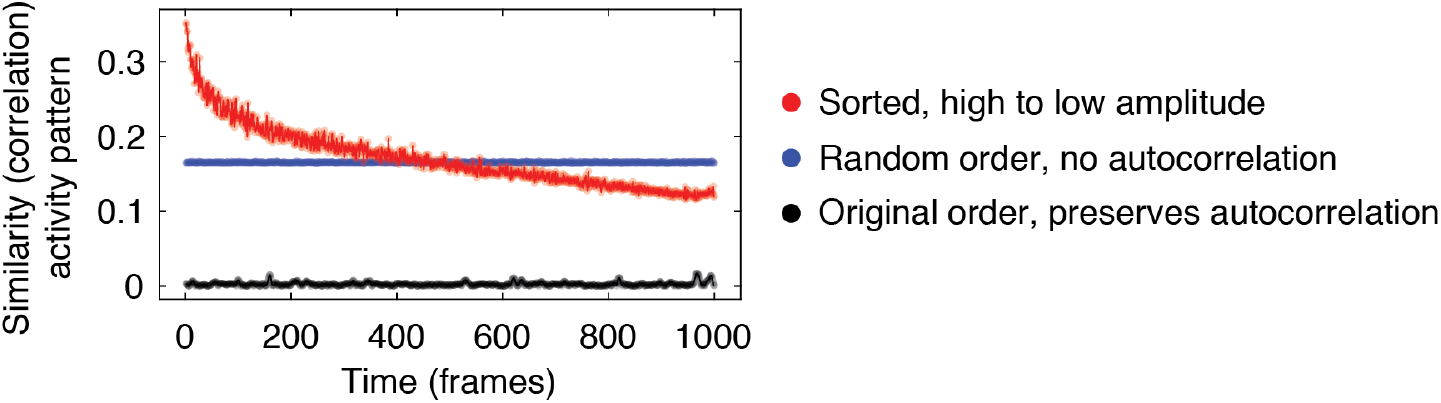
Inter-subject correlations of brain activity during rest. In the main text we demonstrated that “events” were characterized by a shared mode of brain activity. The procedure for doing so involved extracting “event” and “non-event” frames using an arbitrary threshold of 5%. Here, we show similar results by rank-ordering frames. Specifically, we calculate the mean inter-subject similarity (absolute Pearson correlation) of brainwide activity patterns. We do this, first, without reordering the resting-state time series (black curve). Because subjects are not locked to a specific stimulus, we find that, on average, subjects’ activity patterns are uncorrelated. The original ordering of the data preserves, at a single-subject level, autocorrelative properties of the fMRI BOLD time series. Next, we destroy this autocorrelation by randomly reordering each subject’s time series and recomputing mean intersubject similarity. While the overall similarity is slightly greater it is still modest. However, when we reorder frames by their overall amplitude (descending order), we find that high-amplitude frames tend to be more strongly correlated with one another, complementing results in the main text and suggesting that high-amplitude events are underpinnned by a shared pattern of brain activity.

**Table S1.** Movies included in each movie scan.

| Scan | Title | Genre | Runtime |
| --- | --- | --- | --- |
| 1 | Man Up and Go | documentary/emotional | 4m20s |
| 1 | The First 70 | documentary | 3m |
| 1 | Fixation | documentary/adventure | 1m42s |
| 1 | The Living | drama | 2m |
| 1 | SAMSARA | documentary/“unparalleled sensory experience” | 1m35s |
| 1 | Blood Brother | documentary | 2m20s |
| 2 | Birdmen | documentary/adventure | 3m59s |
| 2 | Groomed | drama | 1m30s |
| 2 | Cold | outdoor/sports | 2m |
| 2 | Sleepwalkers | drama | 2m |
| 2 | A Kind of Show | comedy | 1m |
| 3 | Geofish | documentary/adventure | 4m40s |
| 3 | The Debut | outdoor/sports | 3m23s |
| 3 | Dreams of a Life | documentary/mystery | 2m10s |
| 3 | The Front Man | documentary | 2m30s |
| 3 | This Is Vanity | drama | 1m |
| 4 | Planetary | documentary | 4m30s |
| 4 | Sign Painters | documentary | 2m50s |
| 4 | Florida Man | documentary/drama | 2m |
| 4 | The Sleeping Bear | drama | 3m40 |

